# OsMORF protein selectively promotes the dual-localized RNA editing factor OsPGL1 binding to its target RNA

**DOI:** 10.1101/2024.11.08.622588

**Authors:** Haijun Xiao, Qiuren Wang, Yingru Lin, Yang Zhong, Chilin Ma, Honghui Lin

**Affiliations:** Key Laboratory of Bio-Resource and Eco-Environment of Ministry of Education, College of Life Sciences; State Key Laboratory of Hydraulics and Mountain River Engineering, Sichuan University, Chengdu 610065, China

**Keywords:** RNA editing, MORF, PPR, Complex

## Abstract

Post-transcriptional RNA editing plays a key regulatory role and serves as a genetic information proofreading mechanism. In flowering plants, RNA editing typically involves the removal of amino groups from cytidine to produce uridine in both plastids and mitochondria. In our previous work, we identified OsPGL1 as a rare organelle editing factor that regulates both mitochondrial and chloroplast RNA in rice. This protein mediates RNA editing and is associated with several OsMORFs. We speculated that they may form editing complex to regulate editing events, but we are still unclear about the detailed mechanism between OsMORF proteins and OsPGL1, as well as their roles in target RNA recognition and binding. In this study, we found that three OsMORFs can significantly promote the binding of OsPGL1 to target RNA in vitro. Notably, OsMORF8 can promote OsPGL1 to form a homologous dimer, thereby enhancing RNA binding to a greater extent. The E and DYW motifs do not affect the role of MORF proteins in RNA editing events, ruling out the potential influence of these two special motifs. In addition, we predicted a series of supercomplex structures of OsPGL1, the OsPGL1-RNA ligand complex, the OsPGL1-OsMORF2-OsMORF8-OsMORF9 complex, and the OsPGL1-OsMORF2-OsMORF8-OsMORF9-RNA complex. We determined that the conformational changes and newly added positively charged surface caused by OsMORF on OsPGL1 are the main reasons for MORF to promote the binding of OsPGL1 to RNA. Based on some new discoveries, we have expanded the RNA editing complex model, which greatly enhances our understanding of plant RNA editing complexes.

**Short summary:** PPRs and MORFs are two kinds of RNA editing factors, both can mediate organelle RNA editing, the difference between them is that PPR can directly bind target RNA, MORF does not directly bind RNA, but can independently or assist PPR in performing RNA editing. Function of MORF protein in post-transcriptional processing is not clear, our study verifies the function of MORFs in the process of PPR binding RNA through molecular biology. We also analyzed the interaction network between PPR-MORF-RNA from the perspective of structural biology.

## Introduction

The photosynthesis of green plants achieves energy conversion in nature, maintains the carbon-oxygen balance in the atmosphere, and provides a source of energy for the diverse biosphere. In flowering plants, RNA editing occurs in both plastids and mitochondria, playing a crucial regulatory role after transcription and deeply affecting the growth and development of chloroplasts. Therefore, RNA editing has a significant impact on photosynthesis and is essential for the growth and development of plants. RNA editing refers to the process of adding, replacing, and deleting nucleotides at the RNA level to correct mutations that occur at the DNA level. It is a mechanism for correcting genetic information. The term “RNA editing” was first proposed in 1986 to describe the addition and deletion of uridine nucleotides in Trypanosoma mitochondria(Benne et al., 1986). In 1989, C-to-U substitutions were discovered in mitochondrial transcripts of evening primrose, representing the first plant RNA editing found by the scientific community(Hiesel et al., 1989). Currently, we know that A-to-I substitution is the primary form of RNA editing in animals, while C-to-U substitution is the primary form of RNA editing in plants. RNA editing in plant organelles is an especially complex process. Different types of proteins encoded by nuclear genes assemble into an editing complex to perform RNA editing.

To date, the RNA editing factors we have discovered include pentatricopeptide repeat proteins (PPRs), multicellular RNA editing factor proteins (MORFs), also known as RNA editing factor interaction proteins (RIPs), organellar RNA recognition motif proteins (ORRMs), and organellar zinc finger proteins (OZs). The researchers refer to those classes of proteins other than PPR as non-PPR-type RNA editing factors(Sun et al., 2016). PPR proteins are a class of proteins that contain about 35 amino acid repeat sequences. They can be divided into P and PLS subfamilies according to different PPR motifs. The P subfamily only contains the P motif, which is 35 amino acids long, while the PLS subfamily has both the P motif and the L motif, which is 35 or 36 amino acids long, and the S motif, which is 31 amino acids long. According to the sequence characteristics of the PLS subfamily, it can be divided into three subgroups: E, E+ and DYW. PPR proteins located in rice plastids include

OsPPR4(Wang et al., 2017), OsPPR16(Huang et al., 2020), etc., while those located in rice mitochondria include FLO18(Yu et al., 2021), OsPPR939(Zheng et al., 2021), etc. In addition to being located in chloroplasts and mitochondria, there are also some PPR proteins located in the nucleus, such as rice PPR protein OsNPPR1(Hao et al., 2019). So far, only two rice PPR proteins, OsPGL1 and PPR34, have been found to be dual-located.

MORF is a small protein characterized by a typical conserved MORF-box with several peptide motifs (Sun et al., 2013). Ten types of MORF proteins have been found in Arabidopsis, and seven types of MORF proteins have been found in rice. Among them, MORF2 and MORF9 are located in rice chloroplasts, MORF1 and MORF3 are located in rice mitochondria, and MORF8 is located in both rice chloroplasts and mitochondria (Takenaka et al., 2012). MORF proteins affect the editing efficiency of transcripts in organelles and have a significant impact on the growth and development of plant organelles. In Arabidopsis, MORF proteins are involved in complete editing sites at almost all sites in plastids transcripts and many sites in mitochondrial transcripts(Takenaka *et al*., 2012).Many PPR mutants and MORF mutants shared one or more organelle RNA editing walkout events. The editing levels of all 8 target sites affected in the Arabidopsis mitochondrial RNA editing factor MEF13 loss-of-function mutant were reduced in both *morf3* and *morf8* mutants(Glass et al., 2015). The editing efficiency of the target *nad2-842* site of MEF10 is also reduced in *morf8* mutant (Bentolila et al., 2012). MORF1 shares *cox3-257* and *ccmB-566* with MEF21 and MEF19, respectively (Takenaka et al., 2010; Takenaka *et al*., 2012). MORF3 and SLO2 jointly regulate RNA editing at four different RNA transcript editing sites in mitochondria (Zhu et al., 2012). Interestingly, MORF2 interacting with the Arabidopsis editing factor GUN1 also showed reduced RNA editing efficiency at multiple sites and confers a phenotype similar to *gun1* during retrograde signaling(Zhao et al., 2019). MORF2 and MORF9 are extremely important, and almost all chloroplast editing sites have their presence. A recent study showed that down-regulation of MORF protein expression in peaches(PpMORFs) also affected chloroplast RNA editing, especially PpMORF2 and PpMORF9 can regulate stress responses in plant immunity by modulating RNA editing(Zhang et al., 2022). In addition to influencing RNA editing, MORF2 and MORF9 have also been found to interact with tetrapyrrole biosynthesis enzymes(TBS enzymes) by influencing the tetrapyrrole biosynthesis pathway and thus embryogenesis(Yuan et al., 2022).

The *Ospgl1* mutant was obtained by genome editing technology, which caused the leaves to turn pale green, and the chlorophyll content in the leaves was significantly reduced. The starch granules in the leaves were also reduced and less dense compared to the wild type. Transmission electron microscopy revealed that the structure of the chloroplast was damaged, but the OsPGL1 mutant did not affect the structure of the mitochondria. Genetic analysis and biochemical experiments showed that OsPGL1 belongs to the DYW subfamily of PPR proteins. By using its own PPR motif, OsPGL1 can directly recognize and bind to the *ndhD* transcript in the chloroplast to achieve RNA editing at the *ndhD*-C878 site, which regulates the development of chloroplast and photosynthesis in rice. Interestingly, although OsPGL1 can specifically recognize and bind to the *ccmFc* transcript in the mitochondria to perform RNA editing at the *ccmFc*-543 site, the amino acid sequence before and after editing did not change. This may be because the structure of the mitochondria and the assembly of the respiratory chain complex III in the OsPGL1 mutant did not change.

In this study, we found that OsMORF2, OsMORF8, and OsMORF9 proteins can interact with OsPGL1 and significantly enhance OsPGL1 binding to the target RNA probe. By calculating protein structures, we analyzed the interactions between OsPGL1 and OsMORF2, OsMORF8, and OsMORF9. This finding provides a deeper understanding of the mechanisms and effects of DYW subclass proteins exemplified by MORF proteins and OsPGL1. It also gives us a clearer understanding of PPR protein binding to RNA under the action of MORF.

## Results

### Solubilizing modification and in vitro expression of OsPGL1

Natural PPR protein is difficult to homogeneously express in *E. coli* due to multiple repeat units, and OsPGL1 is no exception, this problem has often been encountered in our previous research species, in order to obtain high expression and well homogeneity of OsPGL1 recombinant. We add some auxiliary amino acids to the 5’ and 3’ ends of the OsPGL1 refered by(Yin et al., 2013): the N-terminal (remove the transmembrane domain) add His-Gln-Thr-Pro-Thr-Pro-Pro-His-Ser-Phe as the cap structure, and the C-terminal conserved DYW residue followed by Asp-Lys-Lys-Ala-Leu-Glu-Ala-Tyr-Ile-Glu-Asp-Ala-Gln as the terminal dissolution helix(Supplemental Figure1). These auxiliary amino acids are derived from a well known PPR protein, PPR10 of *Zea mays*, and its good solubility has been verified in several studies of artificially engineered PPR protein(Shen et al., 2016). The modified OsPG1 is termed as mOsPGL1 (modified OsPGL1), and the hydrophobicity/hydrophilicity prediction is carried out by the EXPASY online prediction tool (Expasy - ProtScale). The results show that the hydrophilicity of the modified OsPGL1 at its 5’ and 3’ ends is significantly stronger than that of the unmodified OsPGL1 (Supplemental Figure 2A,B), and then we fuse mOsPGL1 with GST tags for prokaryotic expression. The results showed that the expression of modified OsPGL1 was significantly higher than that of unmodified OsPGL1 (Supplemental Figure 2C).

### MORF proteins significantly promotes OsPGL1 binding to *ndhD* RNA without affecting binding to *ccmFc* RNA

Previously, we have found that OsPGL1 was interacted with OsMORF2, OsMORF8, and OsMORF9, but it remains unclear how MORF proteins affect OsPGL1-mediated RNA editing. To investigate the effect of MORF on OsPGL1 and RNA editing, we designed RNA electrophoresis mobility experiment to detect the effect of three MORF proteins on the ability of OsPGL1 binding to target RNA. We expressed individually mOsPGL1, co-transformed mOsPGL1 with OsMORF2, mOsPGL1 with OsMORF8, and mOsPGL1 with OsMORF9 in *E. coli*, respectively(Supplemental Figure 3). These proteins are concentrated, dialyzed, and leveled so that the concentration of different recombinant proteins maintains a consistent level. And then incubated with the RNA oligonucleotide *ndhD* and *ccmFc* probes where the two target RNA editing sites are located, respectively. Followed by RNA EMSA assay, we found that when *ndhD* was used as the probe, the electrophoretic migration signal after the addition of co-transformed OsPGL1 and three MORRF proteins was stronger than OsPGL1 alone, and the migration signal gradually increased with the increase of co-expression concentration(Figure 1B-D). Indicating that all three MORF proteins could significantly promote OsPGL1 binding to chloroplast target *ndhD*. Moreover, we also found that in the presence of MORF8, there exist two migration signals(Figure 1C), and we hypothesized MORF8 may promote the formation of homodimers of OsPGL1 to bind to target RNA.

**Figure 1.**
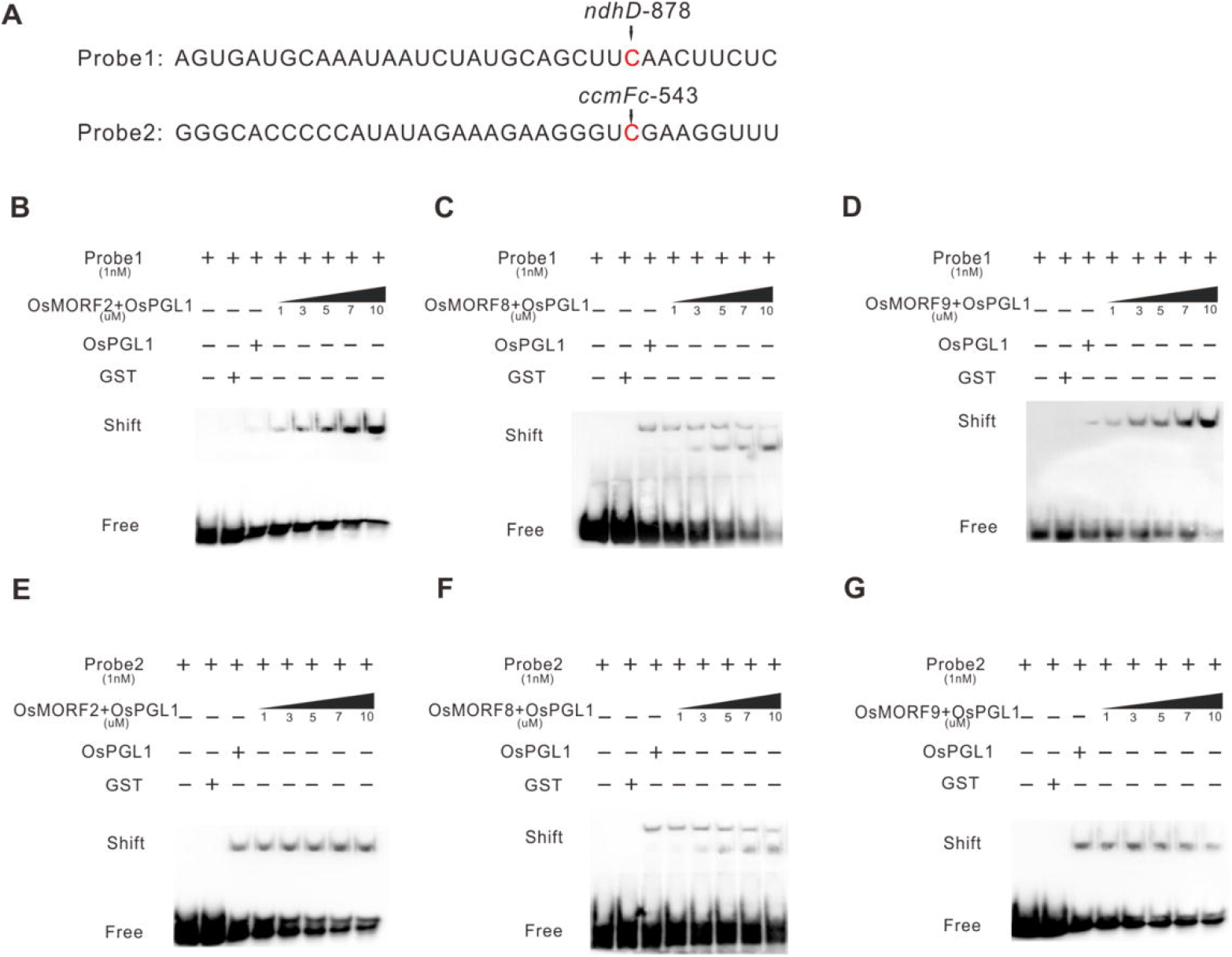
OsMORFs protein can effectively promote the binding of OsPGL1 to its target RNA in different degree. **(A)** Schematic sequences of two OsPG1-bound RNA probes containing target editing sites **(B-D)** Analysis of binding activity of the OsPGL1-OsMORF co-expression substrate to probe 1,GST was used as negative control. **(E-G)** Analysis of binding activity of the OsPGL1-OsMORF co-expression substrate to probe 2,GST was used as negative control.

However, in the components with *ccmFc* target RNA as the probe, when MORF protein was added, the gel migration signal did not have a significant enhancement trend, indicating that MORF protein could not promote the binding of OsPGL1 to the mitochondrial target RNA *ccmFc*(Figure 1E-G). A similar phenomenon is that MORF8 also has the effect of promoting the formation of homodimers of OsPGl1 to bind to target RNA(Figure 1F), This may be because MORF8 is the only dual-localized protein in the MORF family identified thus far. In a word, our results suggest that the MORF protein interacting with OsPGL1 is more inclined to promote the regulation of chloroplast transcripts rather than mitochondria by OsPGL1.

### MORF proteins have distinctive structural domains

MORF proteins are kinds of chaperone proteins that do not interact directly with RNA. These proteins have a conserved domain about 100 amino acids in length called MORF-box. We calculated the protein structures of the MORF family by AlphaFold2, Based on the protein structure(Figure 2A), we used the online program to map the secondary structure topology of MORF. To ensure the accuracy of the topological image, we also performed manual checking. We found that the MORF proteins discovered thus far all have a highly similar secondary structure. Starting from the N-terminal end, they are β-fold, α-helix, α-helix, β-fold, β-fold, α-helix, β-fold, β-fold, and β-fold, respectively. These nine contiguous protein secondary structures are common to all MORF proteins, which contain three α-helices and six β-folds(Figure 2B). These secondary structures act closely together to form a sphere-like surface structure. We now clearly define MORF structural domains as conserved structural domains containing the above secondary structures and call them uniformly MORF-box. We also found that one side of the MORF-box structure has a negative charge distribution on the surface(Figure 2C).

**Figure 2.**
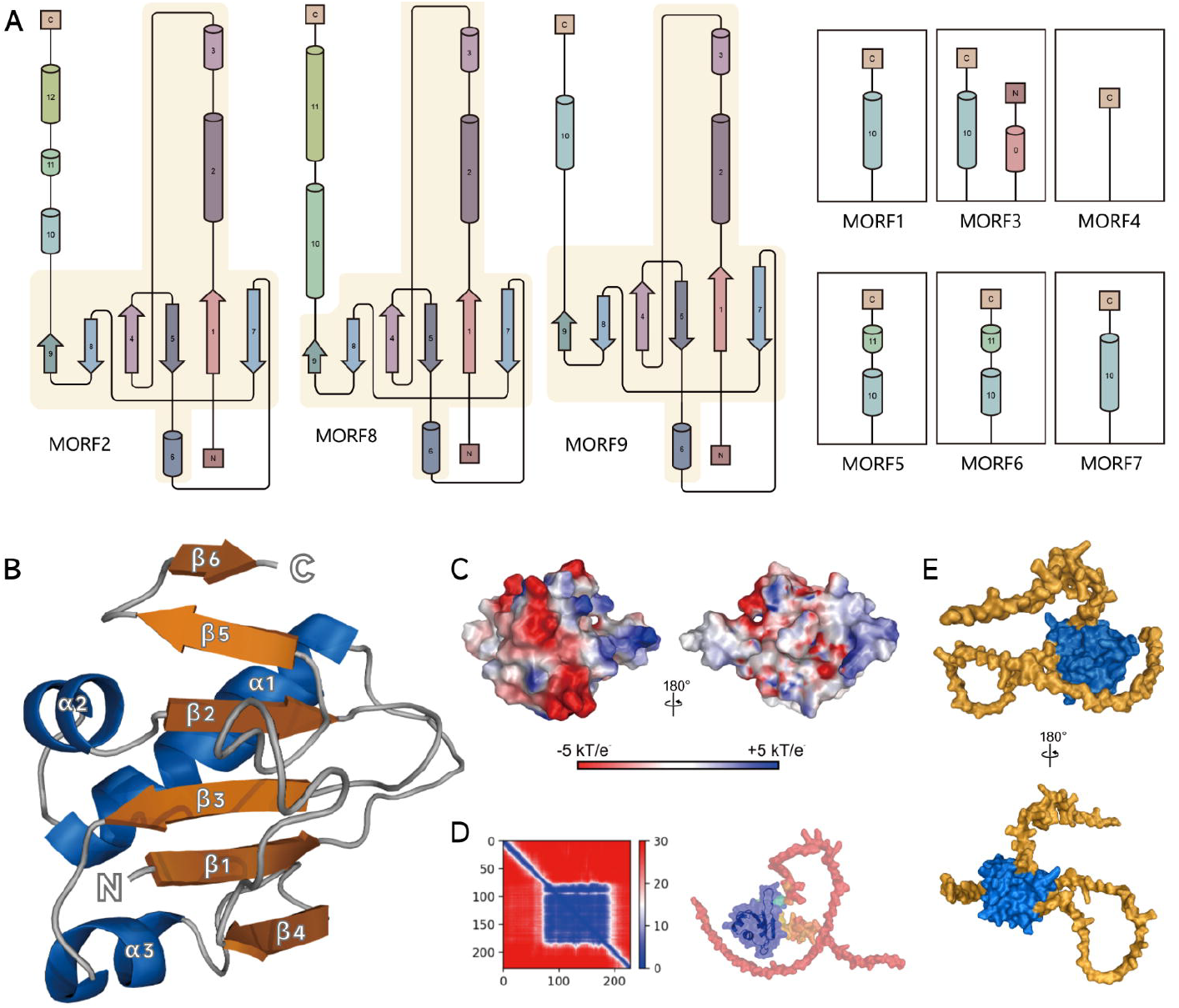
MORFs protein structure. **(A)** Image of the secondary structure topology of the MORF. All secondary structures from the N-terminal to the C-terminal are labeled. There are shades of the MORF-box structural domain common to all MORF proteins. Except for the MORF-box structural domain, all other secondary structures on the protein are indicated separately. **(B)** MORF-box structural domain image. 9 secondary structures, forming a βααββαβββ sequential structural domain. **(C)** Electrostatic surface of MORF9. The electrostatic surface potential of MORF9 protein was obtained by APBS. **(D)** Calculates the score of the result. Blue represents a high confidence level and red represents a low confidence level. **(E)** The surface of MORF protein. The blue is the MORF-box structural domain and the yellow is the free peptide chain.

Previously, the structures of MORF1 and MORF9 were obtained by X-ray diffraction(Haag et al., 2017; Yan et al., 2017). However, suffered to the limitation of the method, only the core structure mainly of MORF-box was obtained. In fact, in addition to the unique MORF-box structure, MORF proteins have extra-long free peptide chains located at both ends(Figure 2E). We found that the number of amino acids in these free peptide chains accounts for more than two-thirds of the total amino acids in the MORF protein. The functions of these free peptide chains will be described later. All calculated protein structures have a high degree of accuracy, with excellent accuracy for the core structure dominated by the MORF-box. The extra-long free peptide chain located at both ends received a lower score, but this is normal; the structure of the free peptide chain is uncertain in vivo(Figure 2D). Since it does not form a secondary structure, it changes with the waves in the liquid environment.

### Prediction of OsPGL1 secondary structure

In our previous study, we have determined that OsPGL1 belongs to the DYW subclass protein in of the PLS subfamily. PLS motif arrangement usually follows the (P1-L1-S1) n-P2-L2-S2 pattern. DYW subclass proteins also have E motifs and DYW motifs. Observation of the OsPGL1 protein structure revealed that the OsPGL1 protein structure as a whole has a right-handed double-steered superhelix structure. (Figure 3A,3B). Its N-terminal consists of 1 separate S-motif and 3 groups P1-L1-S1, of which the third group is a P-L2-S2 triplet structure. The PLS triplet domain is followed by E1, E2 motifs, and DYW motifs. (Figure 3C). The P, L, S, and E motifs consist of two reverse parallel α-helices, which together form the inner and outer sides of the superhelix structure. Starting from the N terminal, the first helix is defined as helix a. Helix a is the inner face of the superhelical structure. The other helix is defined as helix b, and helix b is the outside of the superhelix structure. Helix a is connected to helix b by an ultrashort turn consisting of two residues(Figure 3C,3D). Three groups of P-L-S triplet structures are stacked on top of each other to form a right-handed supercoil. Measurement of the OsPGL1 protein yields a protein polar axis of approximately 129 Å, an equatorial diameter of approximately 58 Å, and an inner diameter of the central cavity of the superhelix structure of approximately 27 Å(Figure 3A).

**Figure 3.**
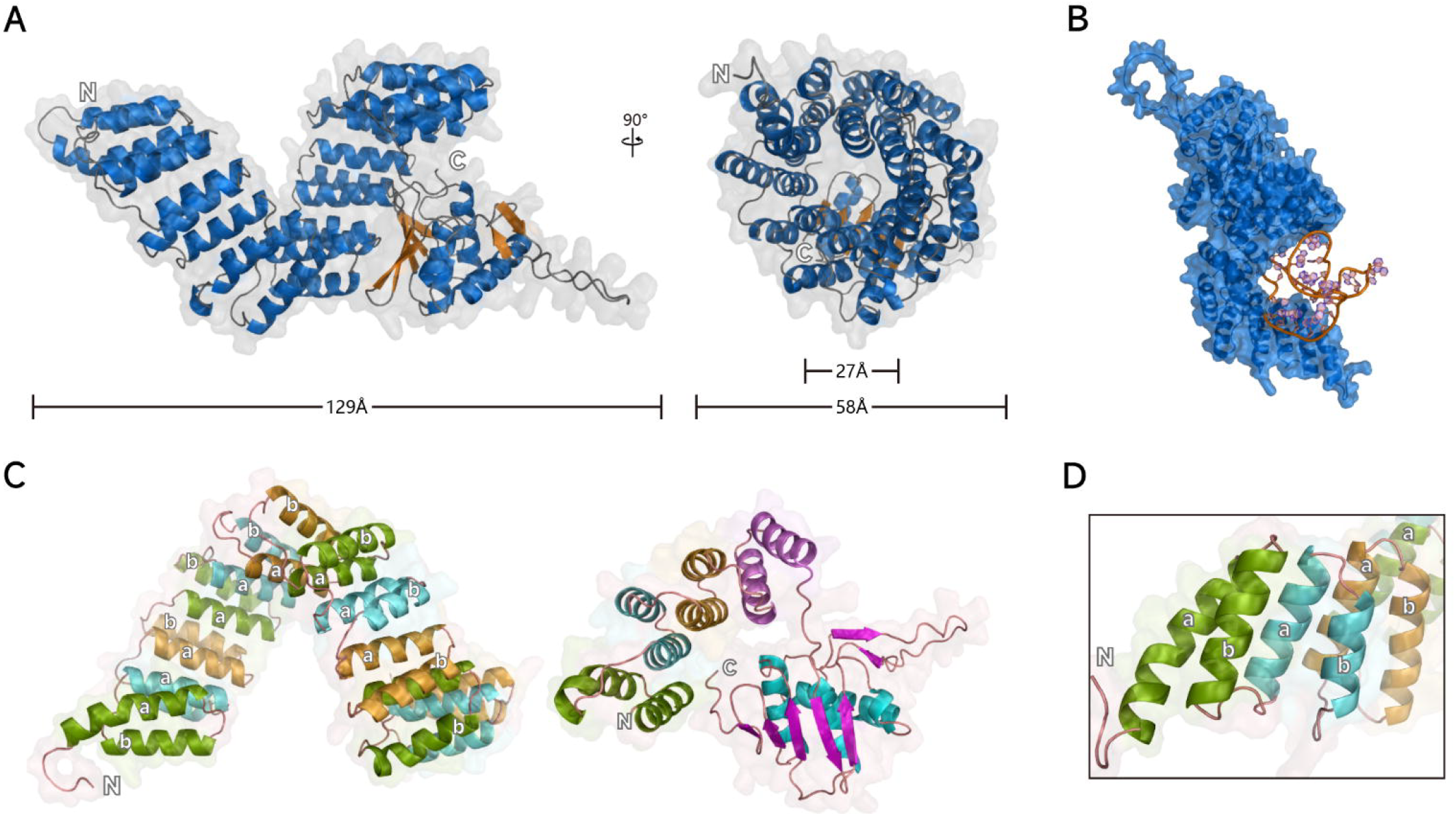
OsPGL1 protein structure. **(A)** Cartoon structure of the OsPGL1 protein. The structure as a whole is a right-handed superhelix. The measured length data is marked on it. **(B)** Image of the OsPGL1 when it binds RNA. The orange RNA is located on the concave side of the superhelix, but is not completely wrapped. **(C)** The picture on the left is the structure of 4 groups of PLS triplets, and the picture on the right is the structure of a group of PLS motifs, E motifs and DYW motifs. Green is the P motif, blue is the L motif, yellow is the S motif, purple is the E motif, and the one on the far right is the DYW motif. **(D)** Enlarged PLS triplet structure. Helix a is the inner side of the superhelical structure and spiral b is the outer side of the superhelical structure.

The secondary structure of the OsPGL1 DYW domain was arranged as β folded-β folded-α helix-β folded-β folded-α helix-β folded-α helix-β folded-β folded-β folded(Supplemental figure 4). As a whole, three α helices are sandwiched between eight β folds. β1, β2, β5, β6, β7, β8 are in the same direction as α2 and α3, and α1 is in the same direction as β3 and β4. Considering the secondary structures in the same direction as a global, the two parts are perpendicular to each other. We speculate that these two parts exercise different roles.

### OsPGL1 attracts targeted *ndhD* RNA through electrostatic interactions

The interaction of the RNA probe with OsPGL1 protein occurs at the PLS motif of the protein. We analyzed the interaction forces between the protein and RNA probe, but unfortunately, we did not find any hydrogen bond, π–π interaction, or cation-pi interaction present between them. Previously, hydrogen bonds between protein and RNA were discovered and the basis for modular recognition of RNA by PPR proteins was found. We obtained the electrostatic surface potential of the protein by APBS and labeled it(Figure 4A-C). We found that the surface of the protein in contact with the RNA probe has a positive charge, which is the fundamental reason for attracting the RNA probe. It is well known that RNA is negatively charged and has an isoelectric point of around 4 due to the exposure of phosphate groups to the molecular surface. Looking at the protein as a whole, the inner side of the PLS motif structure is mostly positively charged and the outer side is mostly negatively charged. α3 and α5 of the PLS motif structure play the largest role(Figure 4D), and the positive charge on their surface is the main reason for the interaction with RNA. The inner side of the PLS motif structure provides some positive charge to interact with the RNA probe by electrostatic forces.

**Figure 4.**
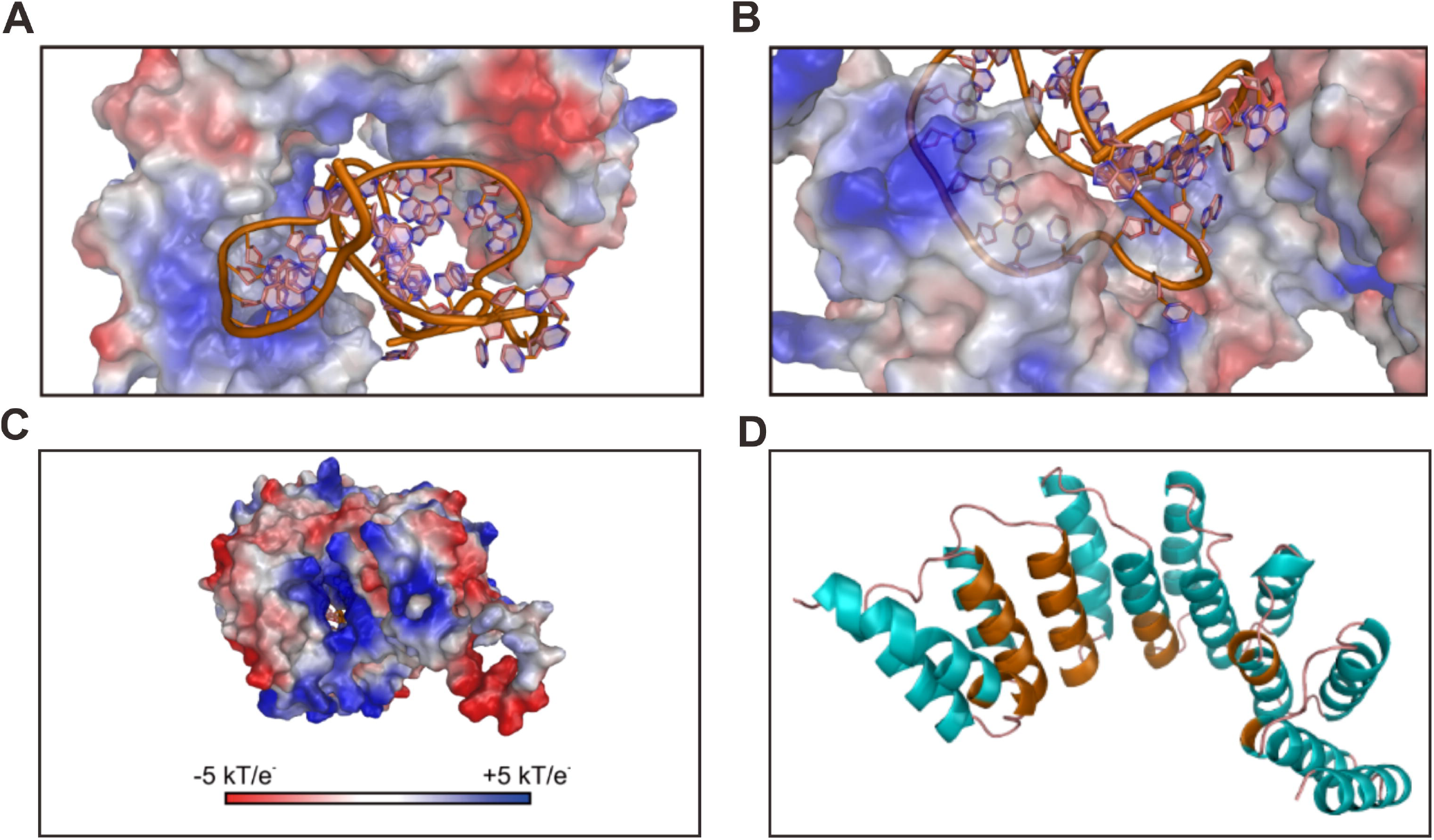
Interaction prediction of OsPGL1 protein with *ndhD* RNA. **(A, B)** The electrostatic surface at the binding site. The electrostatic surface potential of OsPGL1 protein is obtained by APBS. The inner side of the PLS motif structure provides a certain positive charge to interact with the RNA probe by electrostatic forces. **(C)** Electrostatic surface of OsPGL1 protein. The electrostatic surface potential of the OsPGL1 protein was obtained by APBS. There is a small hole near the structural domain of DYW. This small pore is surrounded by a lot of positive charges. We speculate that the RNA strand will pass through here and enter the DYW structural domain. **(D)** The binding site of RNA. The portion of OsPGL1 that contacts RNA is marked in orange. α3 and α5 of the PLS motif structure are the most contacted.

### MORF alters the morphology of OsPGL1 protein to make OsPGL1 bind RNA better

Because OsPGL1 regulates target RNA editing by interacting with three MORF proteins, we simulated possible complex structures between them. We found that the MORF protein caused a striking change in the morphology of OsPGL1. We measured that the polar axis of the OsPGL1 protein in the supercomplex is about 132 A, which is an increase of 3 A compared to the OsPGL1 protein monomer(Figure 5A). The first set of PLS triplet structures starting from the N terminus are positioned closer to the inner side, making the superhelical structure less cylindrical. This would also allow OsPGL1 to bind RNA well.

**Figure 5.**
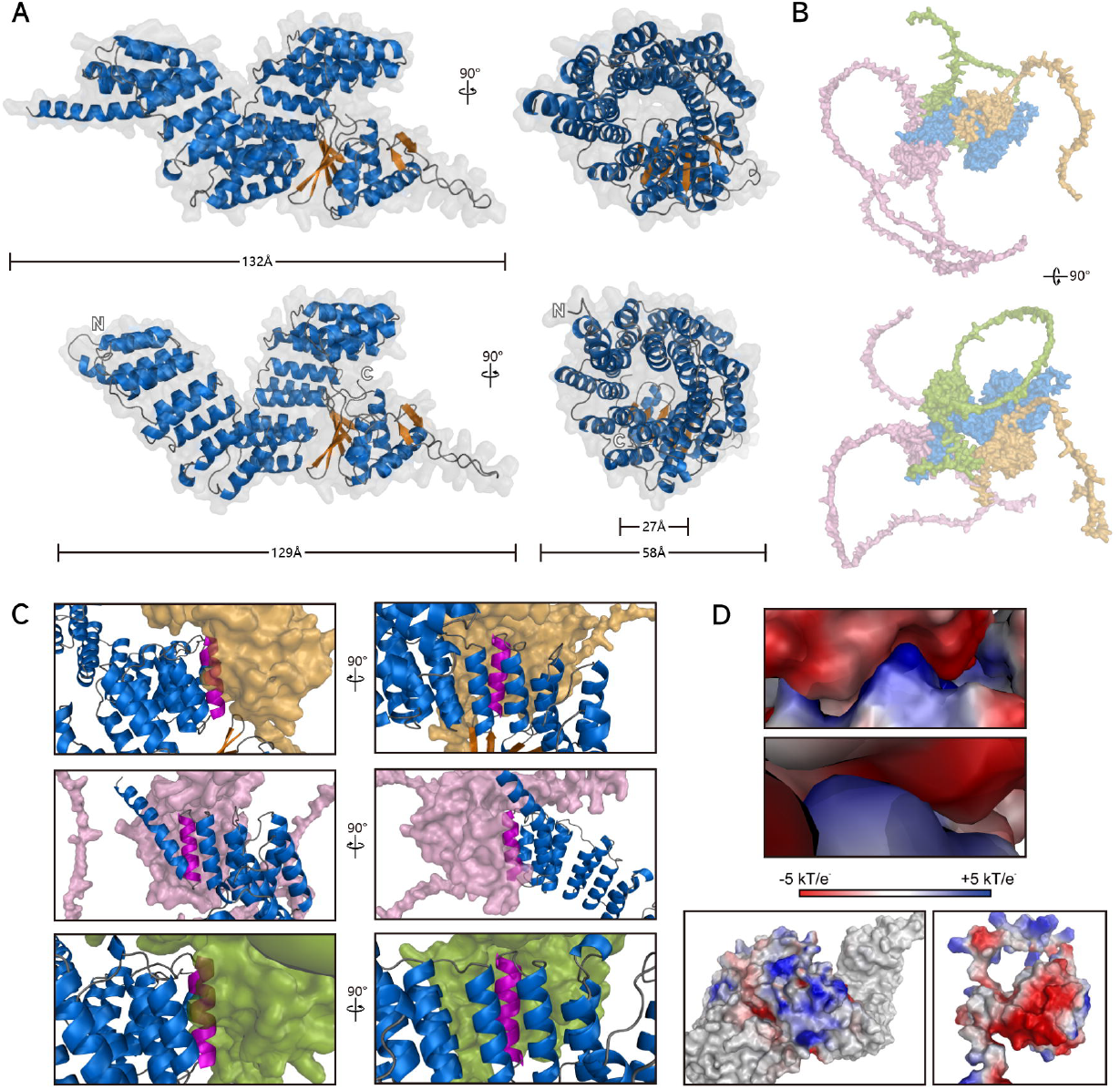
Simulation of the OsPGL1-OsMORFs RNA editing complex. **(A)** The protein located at the top is OsPGL1 in the complex, and the protein located at the bottom is OsPGL1 alone. **(B)** The structure of the supercomplex. The free peptide chain is derived from the MORF protein. **(C)** MORF mainly interacts with the P motif b on OsPGL1. The purple are P pattern b, the orange is OsMORF2, the pink is OsMORF8, and the green is OsMORF9. **(D)** Electrostatic surface of OsPGL1 and MORF binding sites. The lower left protein is OsPGL1, the lower right protein is MORF, the upper one is MORF and the lower one is OsPGL1. The electrostatic surface potential was obtained by APBS.

Studying the interaction interface of OsPGL1 with MORF, we can observe that OsMORF2 interacts with L motif b on the second PLS triplet of OsPGL1, OsMORF8 interacts with N-terminal domain b of OsPGL1, and OsMORF9 interacts with L motif b on the first PLS triplet of OsPGL1. In conclusion, MORF mainly interacts with the L motif b on OsPGL1(Figure 5C). This is consistent with previous studies in which L-motif b was found to be the primary binding site(Yan *et al*., 2017). Then, we analyzed the interaction forces between OsPGL1 and MORF. There are electrostatic interactions, hydrogen bonds, and π-π interactions between them.

Observing the interaction interface between OsMORF2 and OsPGL1(Supplemental Figure 5), it can be observed that both residues VAL-172 and TYR-171 of OsMORF2 are hydrogen-bonded to residue ARG-208 of OsPGL1. In addition, residues ASP-169 and TYR-142 of OsMORF2 both formed hydrogen bonds with the five-membered ring on residue HIS-246 of OsPGL1. Residue TYR-272 of OsPGL1 had hydrogen bonding interactions with residues GLU-158 and VAL-166 of OsMORF2. The above are the important hydrogen bonds between OsMORF2 and OsPGL1, in addition to the π–π interaction between them. The six-membered ring on residue PHE-165 of OsMORF2 has π–π interaction with the five-membered ring on residue HIS-241 and the six-membered ring on residue TYR-272 of OsPGL1.

Then, we observed the interaction interface between OsMORF8 and OsPGL1(Supplemental Figure 5). Residues ASP-167 and TYR-177 of OsMORF8 both form hydrogen bonds with residue LYS-40 of OsPGL1. Residue TYR-169 of OsMORF8 forms a hydrogen bond with residue LYS-37 of OsPGL1. Residue ARG-162 of OsMORF8 and the five-membered ring on residue HIS-68 of OsPGL1 also have hydrogen bonding force. Besides, the five-membered ring on residue HIS-35 of OsPGL1 has π–π interaction with both the five-membered ring and the six-membered ring on residue TRP-163 of OsMORF8.

We also observed the interaction interface of OsMORF9 with OsPGL1(Supplemental Figure 5). Residues ILE-163 and TYR-162 of OsMORF9 both formed hydrogen bonds with residue ARG-138 of OsPGL1. Residues LEU-78, LEU-77 and ILE-76 of OsMORF9 all formed hydrogen bonds with residue GLN-134 of OsPGL1. Residues LEU-78 and LEU-77 of OsMORF9 also have hydrogen bonding force with residue ARG-102 of OsPGL1. The above are the important hydrogen bonds between OsMORF9 and OsPGL1. In addition, the five-membered ring on residue HIS-136 of OsPGL1 has π–π interaction with both the five-membered ring and the six-membered ring on residue TRP-156 of OsMORF9.

Above, we analyzed the important hydrogen bonds and π–π interactions between each of the three MORF proteins and OsPGL1, but as a whole, the important hydrogen bonds and π–π interactions between the MORF proteins and OsPGL1 do not have uniformity. The only thing that can be determined is that the P-mode b in the PLS triplet structure is the main structure that interacts with the MORF protein. Immediately afterward, we obtained the electrostatic surface potentials of the three MORF proteins and OsPGL1 protein by APBS and labeled them(Figure 5D). We are excited by such results. The contact surfaces of all three OsMORF proteins are negatively charged, while all three contact surfaces of OsPGL1 proteins are positively charged, which makes OsMORF proteins and OsPGL1 proteins tightly bound. Probing the camera into the bound protein gap, it was observed that OsMORF protein and OsPGL1 protein bind almost seamlessly due to electrostatic interaction. From this, we conclude that electrostatic interactions are the fundamental reason for the binding of MORF protein and OsPGL1, and only after MORF and OsPGL1 are tightly bound due to electrostatic interactions, hydrogen bonds and π–π interactions are further formed to make the binding of the two proteins stronger.

At the beginning of this section, we have mentioned that the structure of OsPGL1 in the complex has changed surprisingly, and we believe that the structure of the current protein complex is more suitable for RNA due to its RNA tight binding properties, and the RNA in the complex is almost completely closed without much base exposure compared to the OsPGL1 protein alone(Supplemental Figure 6A,C). We still did not find any hydrogen bonding or π–π interactions between RNA and the complex. OsMORF2, OsMORF8, and OsMORF9 all have some interactions with RNA, and these interactions are electrostatic. We obtained the electrostatic surface potential of the protein complex by APBS and labeled it(Supplemental Figure 6B). We compared the OsPGL1 protein alone and found that the positive surface potential of the binding region on the inner side of the complex more than doubled, with OsMORF8 making the largest contribution. Thus, we know that MORF can change the structure of OsPGL1 protein and the electrostatic surface potential of the complex, allowing OsPGL1 to better bind RNA, and it is the positive potential on the surface of the binding region that is critical for RNA binding ability.

## Discussion

In our previous study, we found that OsPGL1 protein can interact with OsMORF2, OsMORF8, and OsMORF9, but we are not sure whether MORF protein can promote OsPGL1 protein to bind target RNA. MORF9 facilitates the artificially designed (PLS)_3_PPR protein to bind RNA(Yan *et al*., 2017). However, the relationship among native PPR proteins, MORF and RNA is poorly understood. The OsPGL1 protein belongs to the DYW subclass of proteins and possesses E and DYW motifs in addition to the PLS motif. Whether these two additional motifs will have an effect on RNA binding is still unknown.

In this study, we explored the effect of MORF protein on the regulation of OsPGL1 protein to its target RNA. We expressed recombinant OsPGL1 and OsMORF2, recombinant OsPGL1 and OsMORF8, and recombinant OsPGL1 and OsMORF9 in *E. coli*. Gel migration results showed that MORF protein could significantly promote OsPGL1 binding to chloroplast transcript *ndhD* RNA. However, there is no obvious promotion effect on the binding of mitochondrial target *ccmFc*. Whether MORF proteins could promote RNA binding to all PLS subfamily proteins needs to be further explored.

To further investigate the basis of OsPGL1 protein-RNA binding, we obtained the OsPGL1-RNA structure. (Figure 3B)Based on the OsPGL1-RNA structure, we were able to determine that the inner side of the superhelical structure formed by multiple PLS triplets on OsPGL1 is the RNA binding site and that electrostatic interactions between the protein and RNA mainly occur. (Figure 4A) Based on the OsPGL1-OsMORF2-OsMORF8-OsMORF9 complex protein structure, we clarified that the interaction between OsPGL1 and MORF is mainly electrostatic(Figure 5B,5D). After binding to MORF, OsPGL1 conformation changes to make it more suitable for binding RNA, and the positively charged surface of the MORF protein also attracts RNA, which structurally explains why MORF can facilitate the binding of OsPGL1 to the target RNA.

In addition to this, we also specifically observed the DYW structural domain(Supplemental Figure 4). The DYW structural domain of OsPGL1 is essentially identical to the structure of OTP86-DYW obtained by X-ray diffraction(Takenaka and Takenaka, 2021). The OTP86-DYW structure has a cytidine deaminase fold and a C-terminal DYW motif with catalytic and structural zinc atoms, respectively. A conserved gating domain within the deaminase fold regulates the active site sterically and mechanically, which is referred to as the gating zinc shuttering process. It is hypothesized that in vivo a certain protein of the plant RNA editor triggers the release of DYW self-inhibition to control and coordinate the cytidine deamination playing a key role in mitochondria and plastids. The activation is triggered by a conformational change in its gating structural domain, which we hypothesize is activated by the RNA bases.

ORRM proteins are also essential for RNA editing, and six ORRM proteins have been identified thus far. ORRM mutants exhibit reduced RNA editing efficiency, but the degree of reduction varies(Hackett et al., 2017). However, as of now, it is not clear what function ORRM proteins perform in the RNA editing complex.

In this study, we reiterate an RNA-editing complex model with OsPGL1 as the core element(Figure 6). PPR proteins serve as their backbone for editing complexes, where PLS motifs are key to identifying target RNA and DYW motifs are where RNA editing actually occurs. where the amine group is removed in the cytidine to produce uridine. The core component of the actual editing is DYW. Previous studies have found that there may be a series of small proteins such as PPO1, OZ, and NUWA around DYW, which constitute the basic components of the RNA editing complex. PPO1 is an enzyme that catalyzes the conversion of protoporphyrinogen IX to protoporphyrin IX in the tetrapyrrole biosynthesis pathway and has been found to play an important role in plastid RNA editing(Zhang et al., 2014). The function of the OZ protein in the RNA editing complex is not known(Sun et al., 2015). The E motif on the PPR protein can recruit a variety of proteins to form an RNA editing core body(Yang et al., 2022). In addition, there are the ORRM proteins, which we speculate are the guide of RNA editing. The ORRM protein binds to the RNA and moves toward the DYW-dominated editing core body, guiding the RNA chain into the editing core body. The function of the MORF protein is unknown, but it has been speculated that it is a bridge between PPR editing factors and deaminase activity in RNA editosome, and that RNA editing efficiency will be greatly improved due to their presence. We know that MORF proteins can interact with PPR proteins, ORRM, PPO1, and OZ1, in other words, MORF proteins can interact with all the RNA editing factors found so far. In addition, the MORF protein has extra-long free peptide chains at both ends. We speculate that the long free peptide chains of MORF proteins could provide a stable space for RNA editing and ensure that RNA editing is not attacked from outside, and the long free peptide chains of MORF proteins could recruit RNA editing factors to participate in a broader range of activities. It is also possible that MORF proteins can act as a bridge between two RNA editing factors, linking the two proteins tightly together.

**Figure 6.**
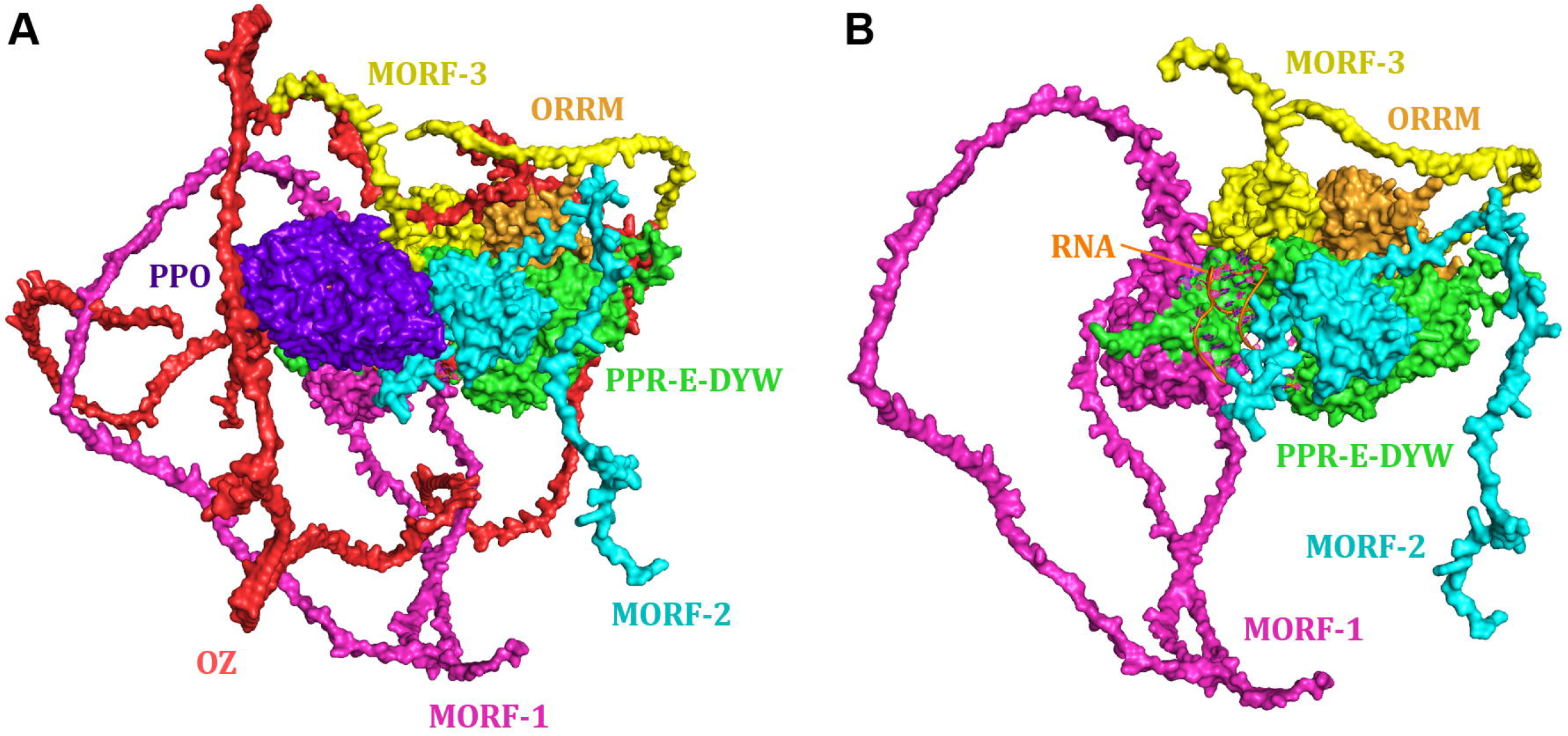
The model of RNA editing complex. The target RNA is obscured by PPO in Figure A and is visible in Figure B.

It has been almost 40 years since the discovery of RNA editing, but we still do not know everything about the RNA editing complex. The research path is a long one for us. However, we can be sure that revealing the composition and function of the plant RNA editing complex will bring great prospects for applications, and further studies on the mechanism of action of RNA editing could lead to a new level of understanding of the life sciences and make RNA editing a powerful new tool for genetic engineering and gene therapy.

## Methods

### Co-transformation of plasmids

The plasmid of OsPGL1 was co-transformed with OsMORFs into competent cells of E. coli BL21 (DE3). After mixing, the cells are ice bathed for 30 min, then heat-stimulated in water at 42 °C for 90 s and then cooled on ice for 2 min The cells were inserted into Luria-Bertani liquid medium containing dual resistance and incubated in a shaker at 37°C for 2 hours. After incubation, the bacterial solution was concentrated to one-third of its original size and applied to Luria-Bertani solid medium containing dual resistance, and incubated overnight. Single colonies were picked for colony PCR, and if both plasmids were confirmed in a single cell, this co-transformation was successful. If the above method is difficult to co-transfer, We transformed the plasmid of OsPGL1 into *E. coli* BL21 (DE3) firstly, and then made the cell into a competent cell. The cells were inserted into Luria-Bertani liquid medium containing resistance and cultured in a shaker at 37°C. When the OD_600_ value of the bacterial solution is about 0.6, collect the cells and bathe the cells in ice for 10 min. Then, collect the cells by high-speed centrifugation and discard the supernatant, and the cells were placed in an ice bath for 10 minutes. Then, the cells were collected by high-speed centrifugation and the supernatant was discarded. Bacteria were resuspended at a ratio of 25 mL of pre-chilled Inoue Buffer per 100 mL of bacterial broth, and the cells were bathed on ice for 10 minutes. After the receptor cells were prepared, the plasmid of OsMORF was first transferred into E. coli BL21 (DE3). Then the incubation and detection were continued according to the first method.

### Protein expression and purification

The cells were inserted into Luria-Bertani liquid medium containing dual resistance and cultured in a shaker at 37°C with 220rpm. When the OD600 value of the bacterial solution was about 0.7, it was induced with 0.5 mM IPTG and induced in a shaker at 37°C with 180rpm for 7 hours. Then, the cells were collected by high-speed centrifugation, and the supernatant was discarded. The cells were then resuspended with Lysis Buffer (PBS and an appropriate amount of lysozyme) and left for 15 minutes. The solution containing the cells was crushed to clarity using an ultrasonic cell disrupter. Cell debris was removed by centrifugation at 12,000 rpm at 4°C. The supernatant was loaded into a column containing Ni^2+^ affinity resin, and the non-target proteins were repeatedly washed with Wash Buffer (PBS and 50 mM imidazole), and the proteins were then eluted with Eluent Buffer (PBS and 250 mM imidazole). The eluted proteins were dialyzed to remove RNAase. After concentrating the proteins, protein purity was checked using sodium dodecyl sulfate polyacrylamide gel electrophoresis (SDS-PAGE) and visualized by Komas Brilliant Blue staining.

### RNA Electrophoretic Mobility Shift Assay

For RNA-EMSA, we synthesized two RNA probes (*ndhD* and *ccmFc*) and used biotin labeling at the 3’ end. The co-expressed proteins were incubated in a 20 μL mixture along with the RNA probes. The mixture was incubated at 25°C for half an hour and then electrophoresed in electrophoresis solution (0.5×TBE). When the bromophenol blue dye enters three-quarters of the gel, electrophoresis is stopped and the reaction is then transferred to a nylon membrane. RNA-EMSA was performed using a kit made by Thermo fisher (Item No.: 20158). The detailed experimental procedures were carried out according to the manufacturer’s instructions.

#### Protein structure prediction and analysis

To better understand the interaction of OsPGL1 protein with MORF proteins, we calculated the structures of OsPGL1 protein, OsMORF2 protein, OsMORF8 protein, and OsMORF9 protein using AlphaFold2. Due to the high confidence of AlphaFold2 prediction results, we could even match some predictions with the structures resolved by the X-ray diffraction method. We first prepare the amino acid sequences of the proteins, then tell AlphaFold2 the amino acid sequences, followed by selecting the computational model and setting additional parameters. The program relies on Google Cloud Server to perform the calculations. After the calculation, the usability of the predictions is evaluated and if the score is above 80, it is fully usable. For specific secondary structures, such as alpha-helix, beta-fold, and beta-turn, AlphaFold2 has a high prediction score. For free peptide chains far from the secondary structure, the scores are lower. However, most of these free peptide chains are also free in the cell(Jumper et al., 2021). After obtaining the structures of the proteins, we analyzed them with a series of tools such as APBS and HDock, and further visualized them on PyMOL.

## Auther contributions

H.X. conceived the project; H.X., Q.W. and Y.L performed the experiments; Q.W. analyzed the protein secondary; H. L. provide valuable suggestions; H.X. wrote the manuscript with feedback from all authors.

