## Supplementary figures and images for "OsMORF protein selectively promotes the dual-localized RNA editing factor OsPGL1 binding to its target RNA"

### Supplemental Figure1

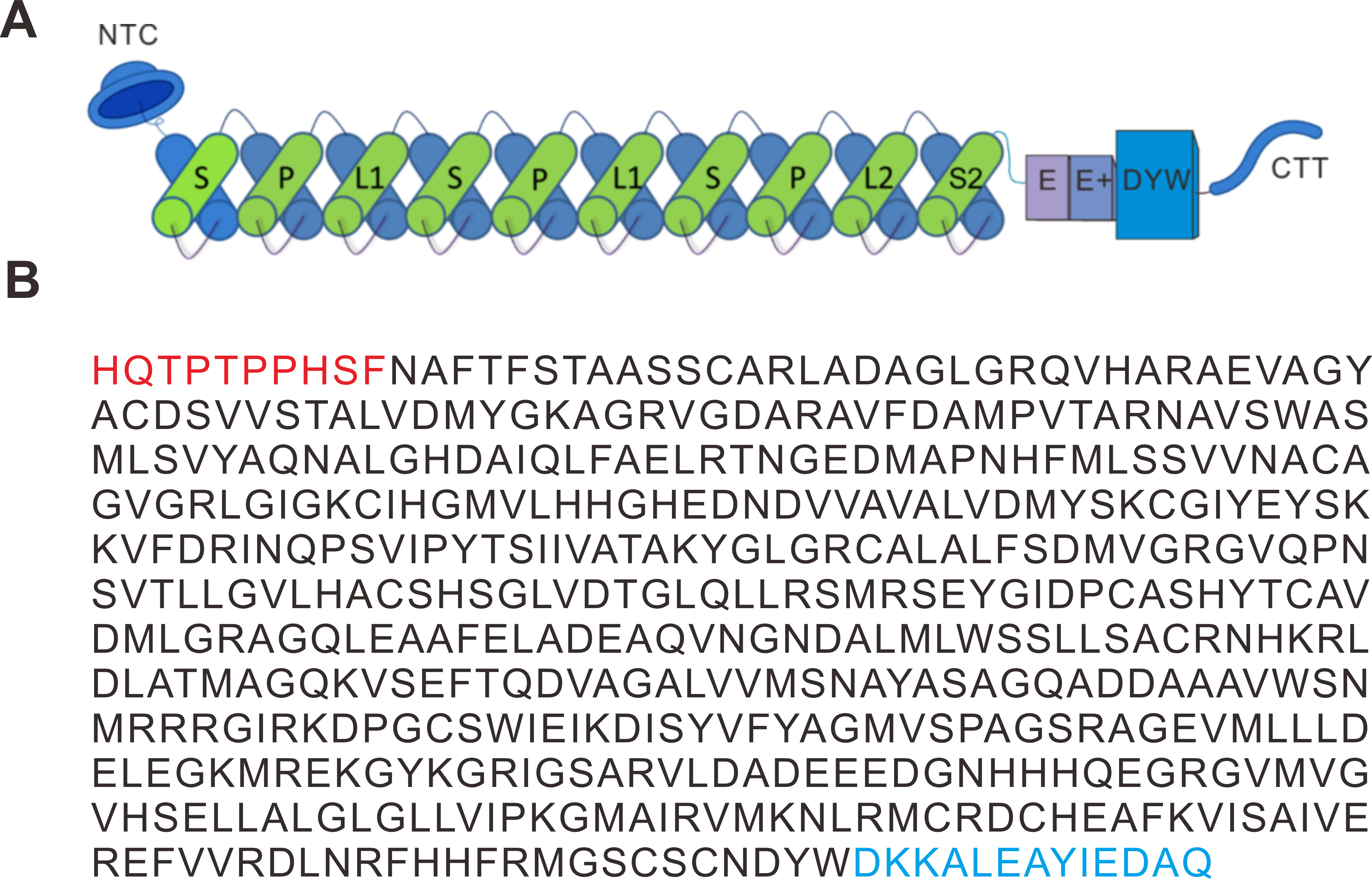

### Supplemental Figure2

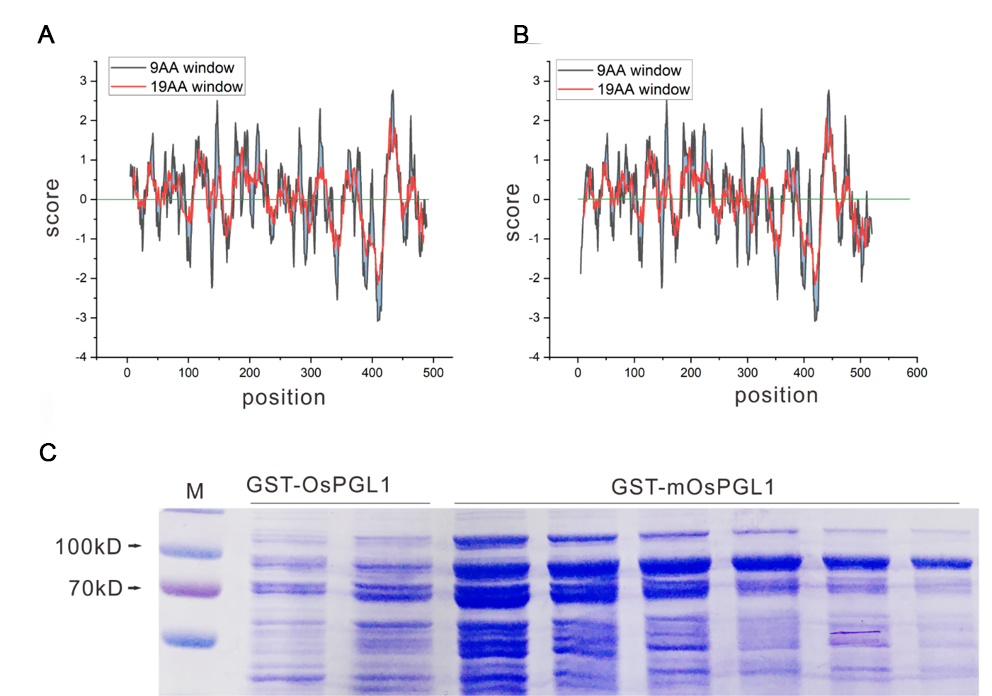

### Supplemental Figure3

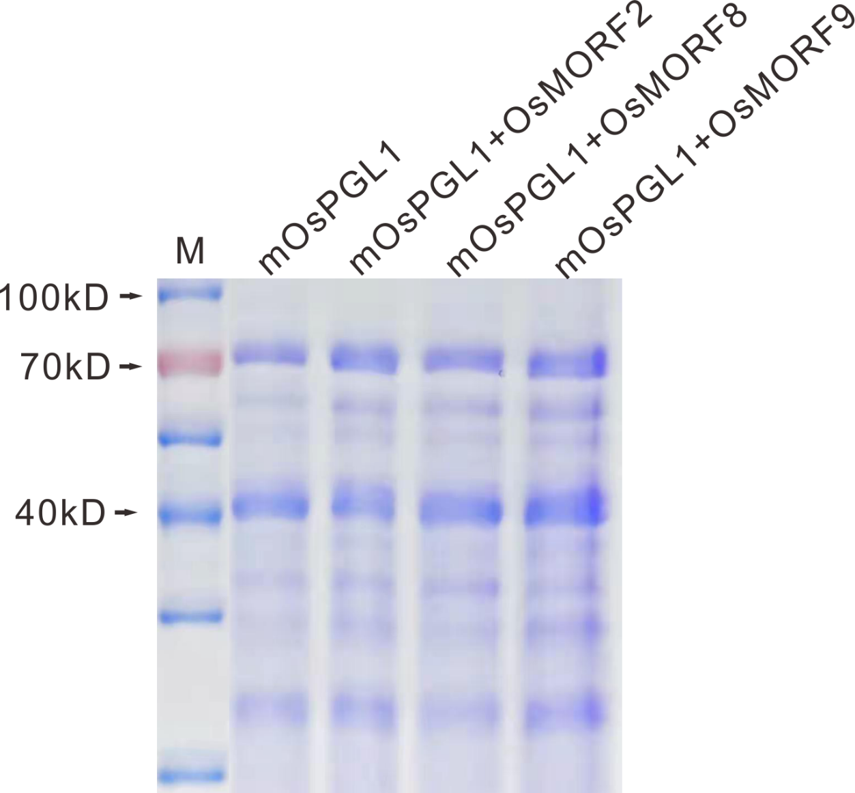

### Supplemental Figure4

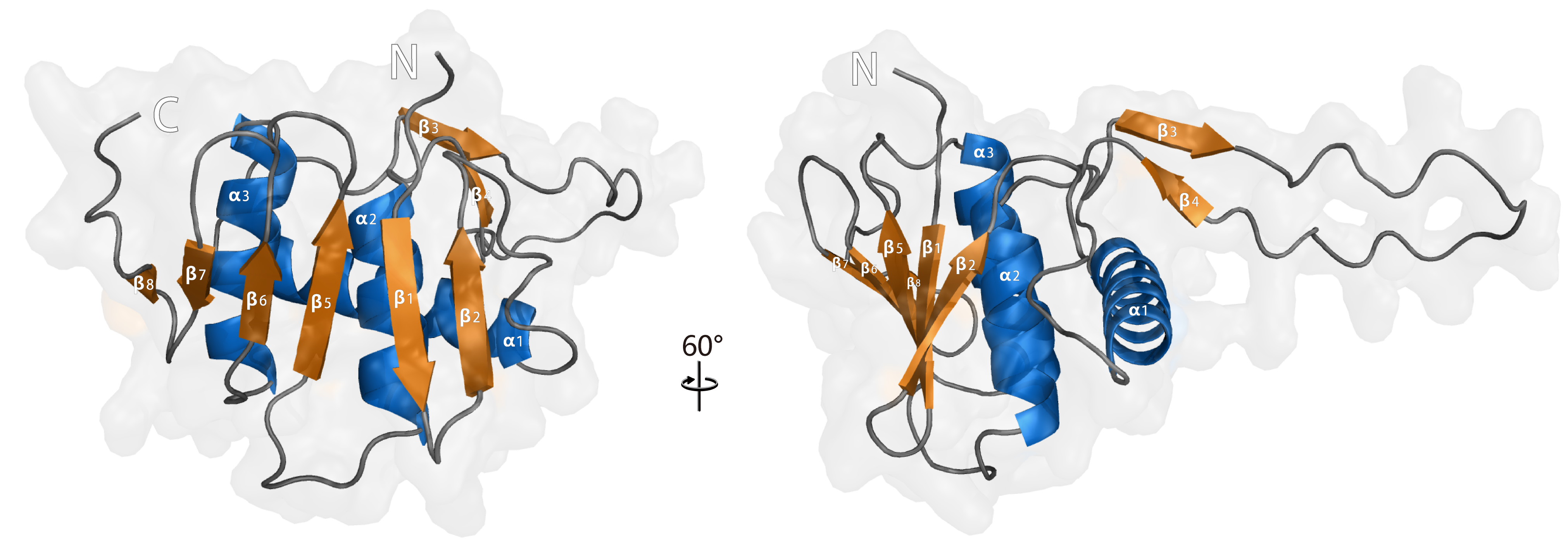

### Supplemental Figure5

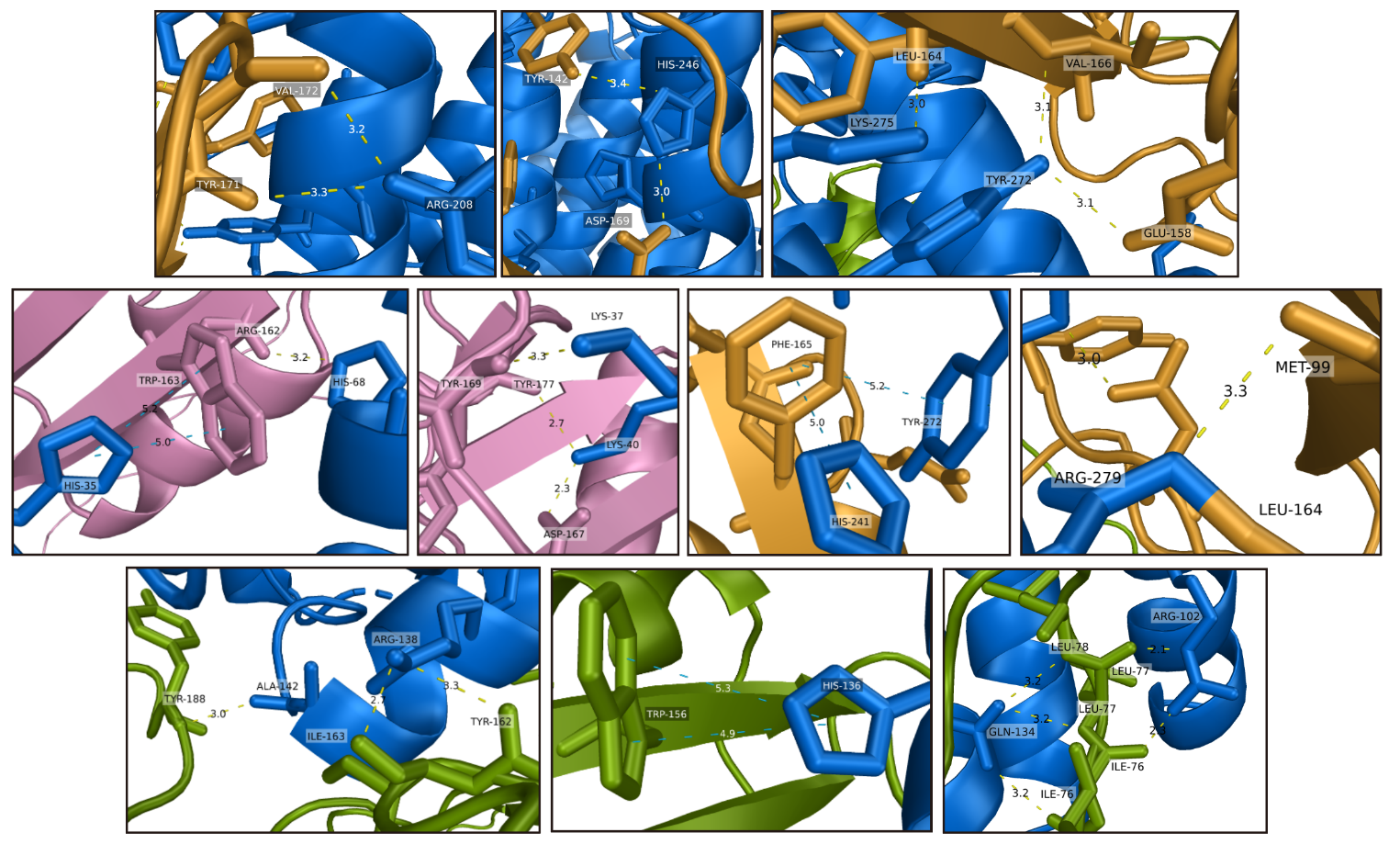

### Supplemental Figure6

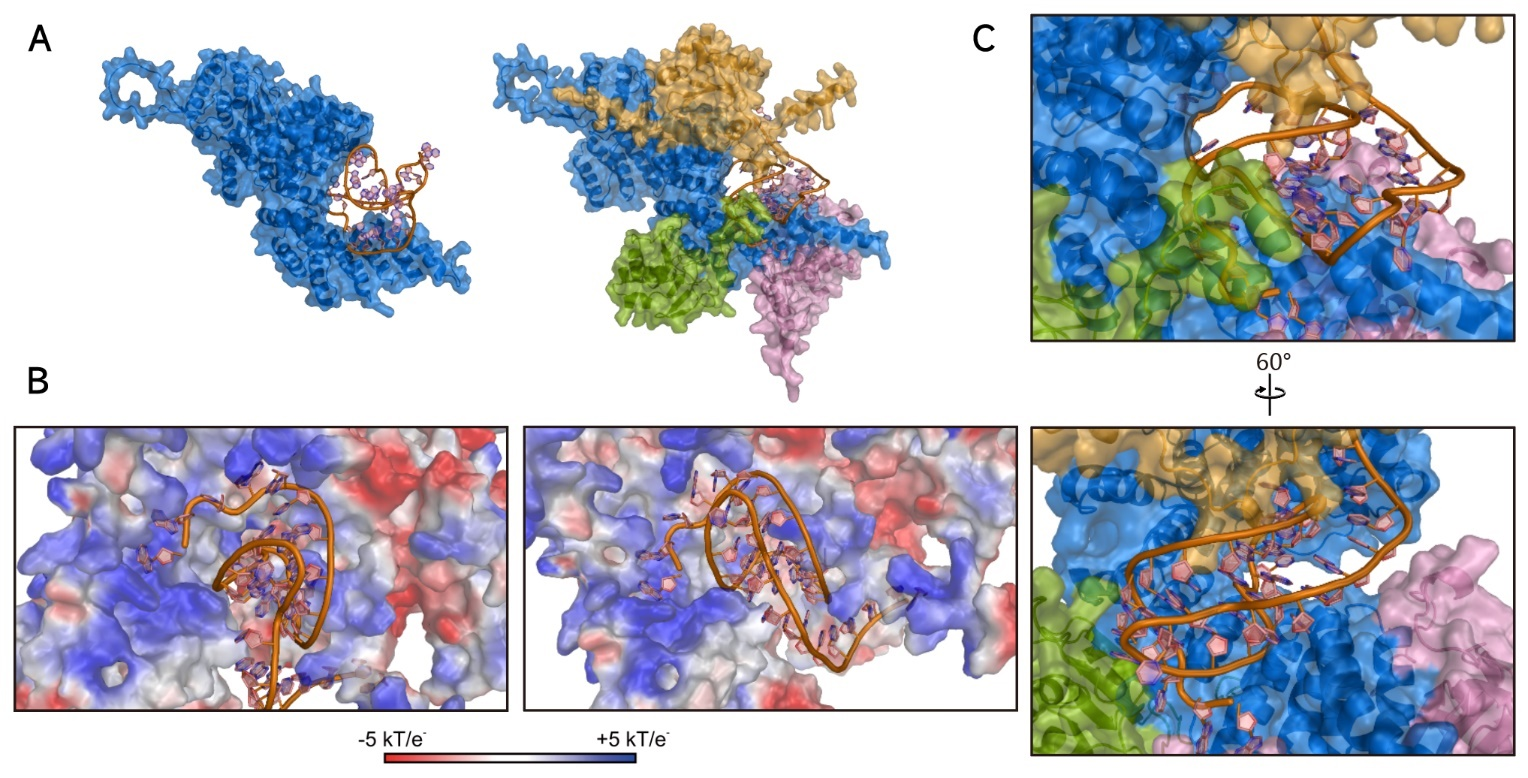
