## Supplemental Table for "OsMORF protein selectively promotes the dual-localized RNA editing factor OsPGL1 binding to its target RNA"

| **Primer name** | **Forward sequence (5’-3’)** | **Reverse sequence (5’-3’)** |
| --- | --- | --- |
| *mOsPGL1*-  EX | AAATCGGATCTGGTTCCGCGTGGATCCCACCAGACGCCGAC | TCGAGAGATCTGTCGACGATATCGAATTCCTGCGCATCCTCTATA |
| *MORF2-* Ex | GGATCCATGGCCGCCGCCGCCGCCGCCGCCG | GGATCCCCGTTGGTTCTCCCTCCTCCGGGC |
| *MORF8-* Ex | GGATCCATGGCGTCGGCGTCGCGCT | GGATCCCTGGTAATTCCTCCCTTGTC |
| *MORF9-* Ex | GGATCCATGGCTTCCTTCCCGACCACC | GGATCCTGAAGAAGAGGCTGACTCAG |
